# Skin transcriptome profiles associated with coat color in goat

**DOI:** 10.1101/028340

**Authors:** Yongdong Peng, Xiaohui Liu, Liying Geng, Chuansheng Zhang, Zhengzhu Liu, Yuanfang Gong, Hongqiang Li, Xianglong Li

## Abstract

Capra hircus, an economically important livestock, plays an indispensable role in the world animal fiber industry. To identify additional genes that may play important roles in coat color regulation, Illumina/Solexa high throughput sequencing technology was used to catalog the global gene expression profiles in the skin of three different coat colors goat (Lubei white goat (white), Jining gray goat (gray) and Jianyang big ear goat (brown)). The RNA-Seq analysis generated 83174342, 70222592 and 52091212 clean reads in white skin, gray skin and brown skin, respectively, which provided abundant data for further analysis. A total of 91 genes were differentially expressed between the gray skin and white skin libraries, with 74 upregulated and 17 genes downregulated. Between the brown skin and white skin libraries, there were 23 upregulated genes and 44 downregulated genes, while there were 33 upregulated genes and 121 downregulated genes between the brown skin and gray skin libraries. To our surprise, *MC1R, MITF, TYR, KIT* and *KITLG* showed no significant difference in the skin of three different coat colors and the expression of *ASIP* was only detected in white skin and not in gray and brown skins. The expression of *PMEL*, *TRPM1*, *DCT*, *TYRP1* and *ELOVL3* was validated by real-time quantitative polymerase chain reaction (qPCR) and the results of the qPCR were consistent with the RNA-seq except the expression of *TYRP1* between the gray skin and white skin libraries. This study provides several candidate genes that may be associated with the development of diferent coat colors goat skin. More importantly, the fact that the *ASIP* gene was only detected in the white skin and not in the other dark skins and the *MC1R* gene showed no significant difference in expression between the three different coat colors goat is of particular interest for future studies that aim to elucidate theirs functional role in the regulation of skin color. These results will expand our understanding of the complex molecular mechanisms of skin physiology and melanogenesis in goat and provide a foundation for future studies.

---

The domestic goat (Capra aegagrus hircus) is a subspecies of goat domesticated from the wild goat of southwest Asia and Eastern Europe. The goat is a member of the family Bovidae and is closely related to the sheep as both are in the goat-antelope subfamily Caprinae. Goats have long been used for their milk, meat, hair and skins throughout the world (Wang *et al*. 2011a; Wang *et al*. 2011b; Xu *et al*. 2013).

Capra hircus, an economically important livestock, plays an indispensable role in the world animal fiber industry. Fiber diameter, length and color are key traits contributing to the economic value of goat and are determined by both genetics (Bunge *et al*. 1996; Lamoreux *et al*. 2001) and other factors, including the environment and certain drugs (Kidson and Fabian 1981; Dereure 2001; Jablonski and Chaplin 2010; Sturm and Duffy 2012). Factors that determine coat color in goat are becoming of increasing interest. White fleece holds greatest economic value due to its ability to be dyed to virtually any color, where as interest in natural colors is increasing due to the green revolution and consumer preference for natural products.

In adult animals, both hair and skin color depend on pigment produced by melanocytes at the base of the epithelium (Cieslak *et al*. 2011). Melanocytes in mammals and birds produce two types of melanin, black to brown eumelanin and yellow to reddish brown pheomelanin (Ito *et al*. 2000; Ito and Wakamatsu 2008). The basic coat colourations are defined by the ratio of the two pigments eumelanin and pheomelanin (Cieslak *et al*. 2011).

In animals, melanic coloration is often genetically determined and associated with various behavioral and physiological traits, suggesting that some genes involved in melanism may have pleiotropic effects (Ducrest *et al*. 2008). At present, a large number of genes have been found to play well-known roles in pigmentation and the analysis of these genes has identified many single nucleotide polymorphisms (SNPs), e.g., *ASIP*, *MC1R*, *TYR, TYRP1*, *DCT*, *MITF*, *KIT, KITLG, OCA2*, *SLC24A4*, *PMEL* (Brunberg *et al*. 2006; Gutierrez-Gil *et al*. 2007; Sulem *et al*. 2007; Deng *et al*. 2009; Nan *et al*. 2009; Duffy *et al*. 2010; Minvielle *et al*. 2010; Nicoloso et al. 2012; Hart et al. 2013; Fan *et al*. 2014; Li *et al*. 2014; Becker *et al*. 2015).

Coat color genes are good candidates for facilitation of trace ability of animal breeds. Several previous studies have paid significant attention to the coat color of animals and showed that this color is determined by the amount and type of melanin produced and released by the melanocytes present in the skin (Ito *et al*. 2000; Ito and Wakamatsu 2008). For example, recent researches have demonstrated that *MC1R* and *ASIP* are known to be major regulators of coat color in mice and *MC1R* and *ASIP* loci are functionally linked to undesirable coat color phenotypes in sheep (Vage *et al*. 1999; Slominski *et al*. 2004; Steingrimsson *et al*. 2006; Norris and Whan 2008). In addition, tyrosinase-related protein 1 (*TYRP1*) is a strong positional candidate gene for color variation in Soay sheep (Gratten *et al*. 2007). Recent studies have combined SNP analysis and gene expression profiling to dissect the basis for the piebald pigmentation phenotype in Merino sheep (Garcia-Gamez *et al*. 2011). It was reported that color variation was likely to stem from differences in the expression levels of genes belonging to the melanocortin in the tawny owl (Emaresi *et al*. 2013). Despite considerable knowledge of the genetic regulation of coat color in some animals, the molecular and cellular mechanisms regulating coat color in fiber-producing species, such as the goat, are not completely understood. This information is critical not only to enhanced basic understanding of regulation of melanogenesis, but also to the identification of novel pharmacological and molecular genetics approaches to regulate or select for coat color in fiber producing species.

RNA-Seq technology is a high-throughput sequencing platform allowing us to detect transcripts with low abundance, identify novel transcript units, and reveal their differential expression between different samples (Wang *et al*. 2009; Wilhelm and Landry 2009; Ozsolak and Milos 2011). To investigate the different expression profiles of the genes involved in goat skin pigmentation and gain insight into molecular mechanisms responsible for biochemistry of skin and fibers (including pigmentation) in animals producing hair, we investigated the transcriptome profiles in skin of goat of different coat colors using high throughput RNA deep sequencing. This will enable us to understand the molecular mechanisms involved in skin pigmentation and provide a valuable theoretical basis for the selection of the excellent natural colors trait.

## MATERIALS AND METHODS

### Ethics Statement

All of the animals were handled in strict accordance with good animal practices as defined by the relevant national and/or local animal welfare bodies. The experimental procedure was approved by the Animal Care and Use Committee of Hebei Normal University of Science and Technology, China and was performed in accordance with the animal welfare and ethics guidelines.

### Goat skin sampling and total RNA extraction

Housing and care of goat and collection of skin samples for use in the described experiments were conducted in accordance with the International Guiding Principles for Biomedical Research Involving Animals (http://www.cioms.ch/frame 1985 texts of guidelines.htm). The animals were locally anaesthetized with hydrochloridum (1.5 ml of 3%, i.h.), following the approval provided by the Animal Hospital of Hebei Normal University of Science and Technology to decrease the animal suffering. 3 Lubei white goats, 3 Jining gray goats and 3 Jianyang big ear goats were selected for sample collection from the goat farm in Laiwu, Shandong province, China. A piece of skin (8 mm in diameter) from the leg was collected via punch skin biopsy under local anesthesia and immediately placed in liquid nitrogen. Total RNA from the sample was extracted using Trizol reagent (Invitrogen, USA) according to the manufacturer’s instructions. The RNA integrity was evaluated by gel electrophoresis and the RNA purity was checked by the ratio of OD260/OD280 and RIN value. RNA samples with RIN value greater than 7.5 and OD260/OD280 ratio greater than 1.75 were selected for goat sequencing.

### Library generation and sequencing

Three RNA samples from white, gray and brown goat skin were pooled before mRNA isolation. Sequencing libraries were generated using NEBNext^®^ Ultra™ RNA Library Prep Kit for Illumina^®^ (NEB, USA) following manufacturer’s recommendations and index codes were added to attribute sequences to each sample. Briefly, mRNA was purified from total RNA using poly-T oligo-attached magnetic beads. Fragmentation was carried out using divalent cations under elevated temperature in NEBNext First Strand Synthesis Reaction Buffer（5X）. First strand cDNA was synthesized using random hexamer primer and M-MuLV Reverse Transcriptase (RNase H^-^). Second strand cDNA synthesis was subsequently performed using DNA Polymerase I and RNase H. Remaining overhangs were converted into blunt ends via exonuclease/polymerase activities. After adenylation of 3’ends of DNA fragments, NEBNext Adaptor with hairpin loop structure were ligated to prepare for hybridization. In order to select cDNA fragments of preferentially 150~200 bp in length, the library fragments were purified with AMPure XP system (Beckman Coulter, Beverly, USA). Then 3 μl USER Enzyme (NEB, USA) was used with size-selected, adaptor-ligated cDNA at 37°C for 15 min followed by 5 min at 95 °C before PCR. Then PCR was performed with Phusion High-Fidelity DNA polymerase, Universal PCR primers and Index (X) Primer. At last, PCR products were purified (AMPure XP system) and library quality was assessed on the Agilent Bioanalyzer 2100 system.

The clustering of the index-coded samples was performed on a cBot Cluster Generation System using TruSeq PE Cluster Kit v3-cBot-HS (Illumia) according to the manufacturer’s instructions. After cluster generation, the library preparations were sequenced on an Illumina Hiseq 2000 platform and 100 bp paired-end reads were generated.

### Reads mapping to the reference genome

Reference genome and gene model annotation files were downloaded from genome website directly. Index of the reference genome was built using Bowtie v2.0.6 and paired-end clean reads were aligned to the reference genome using TopHat v2.0.9. We selected TopHat as the mapping tool for that TopHat can generate a database of splice junctions based on the gene model annotation file and thus a better mapping result than other non-splice mapping tools.

### Differential expression analysis

The expression abundance of each assembled transcript was measured through Reads Per Kilobase of exon model per Million mapped reads (RPKM). Prior to differential gene expression analysis, for each sequenced library, the read counts were adjusted by edgeR program package through one scaling normalized factor. Differential expression analysis of two conditions was performed using the DEGSeq R package (1.12.0). The P values were adjusted using the Benjamini & Hochberg method. Corrected P-value of 0.005 and log2(Fold change) of 1 were set as the threshold for significantly differential expression.

### GO and KEGG enrichment analysis of differentially expressed genes

Gene Ontology (GO) enrichment analysis of differentially expressed genes was implemented by the GOseq R package, in which gene length bias was corrected. GO terms with corrected Pvalue less than 0.05 were considered significantly enriched by differential expressed genes. KEGG is a database resource for understanding high-level functions and utilities of the biological system, such as the cell, the organism and the ecosystem, from molecular-level information, especially large-scale molecular datasets generated by genome sequencing and other high-through put experimental technologies (http://www.genome.jp/kegg/). We used KOBAS software to test the statistical enrichment of differential expression genes in KEGG pathways.

### Real-time quantitative RT-PCR

The expression of these genes was quantified by qRT-PCR using QuantiTect SYBR Green RT-PCR (Qiagen, Waltham, MA). Information regarding the primers of *PMEL*, *TRPM1*, *DCT*, *TYRP1*, *ELOVL3* used for the qPCR can be found in Table 1. *GAPDH* was used as housekeeping gene. Quantitative real-time PCR was performed in triplicate on the Stratagene iQ5 system. The 20 μL PCR reaction included 10 μL SYBR Premix Ex Taq II (TaKaRa, Dalian,China), 0.5 μL specific forward primer, 0.5 μL reverse primer, 0.4 μL ROX reference dye, 0.5 μL diluted cDNA and 8.6 μL RNase free water. Cycling parameters were 95°C for 4 min, followed by 40 cycles of 95°C for 15 sec, 56°C or 58°C for 30 sec and 72°C for 45 sec. Melting curve analyses were performed following amplifications. At the end of the cycles, melting temperatures of the PCR products was determined between 70°C and 90°C. The iQ5 software (Bio-Rad) was used for detection of fluorescent signals and melting temperature calculations. Quantification of selected mRNA transcript abundance was performed using the comparative threshold cycle (CT) method. The difference in abundance of mRNA for the genes was determined by analysis of variance.

**Table 1.**
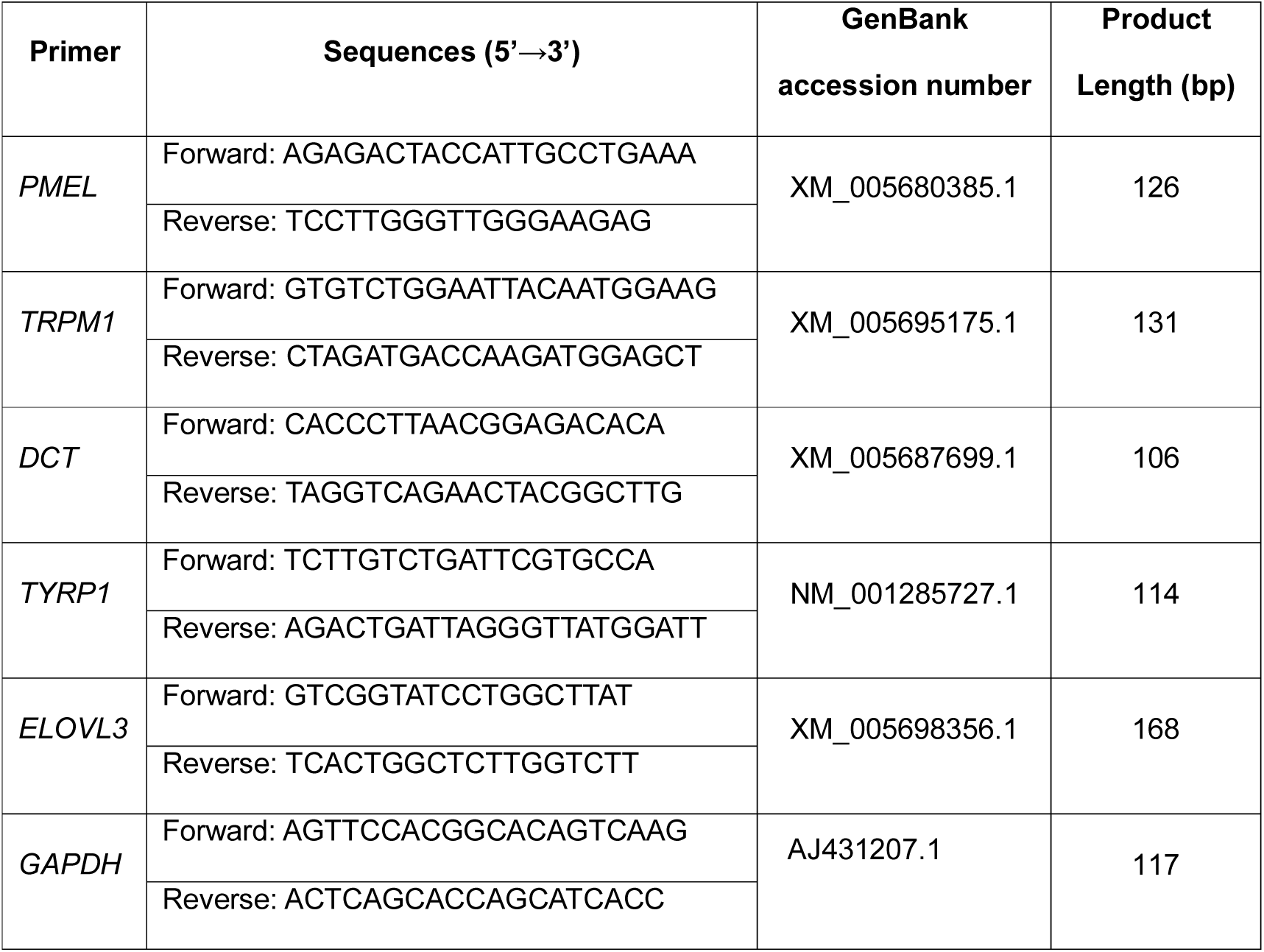
Information regarding the specific primers used for the qPCR.

## DATA AVAILABILITY

Table S1 contains top 30 highly expressed genes in goat skin. TableS2 covers differentially expressed genes between the gray and white skin. Table S3 covers differentially expressed genes between the brown and white skin. Table S4 covers differentially expressed genes between the brown and gray skin. Table S5 contains the list of GO categories for DEGs between different coat colors goat skin. Table S6 encompass KEGG pathway analysis for the gray and white skin differentially expressed genes. Table S7 encompass KEGG pathway analysis for the brown and white skin differentially expressed genes. Table S8 encompass KEGG pathway analysis for the brown and gray skin differentially expressed genes.

## RESULTS

### Massively Sequencing and Aligning to the Reference Genome

To maximise the coverage of the three different coat colors goat (Lubei white goat (white), Jining gray goat (gray) and Jianyang big ear goat (brown)) skin mRNA by RNA sequencing, RNA libraries were constructed by pooling RNA isolated from different coat colors individuals as a sample library. These RNA-Seq libraries generated over 54 million raw reads from each library. After filtering the only adaptor sequences, those containing N sequences and low quality sequences, the three RNA-Seq libraries still generated over 26 million paired-end clean reads in each library, and the percentage of paired-end clean reads among raw tags in each library ranged from 94.07% to 94.71% (Table 2 and Figure 1).

**Table 2.** Quality Control of the Sequencing Data

| Sample name | Raw reads | Clean reads | clean bases | Error rate(%) | Q20(%) | Q30(%) | GC content(%) |
| --- | --- | --- | --- | --- | --- | --- | --- |
| White_1 | 44207310 | 41587171 | 5.2G | 0.04 | 94.95 | 89.92 | 51.36 |
| White_2 | 44207310 | 41587171 | 5.2G | 0.04 | 93.45 | 87.66 | 51.35 |
| Gray_1 | 37218908 | 35111296 | 4.39G | 0.04 | 94.79 | 89.62 | 51.39 |
| Gray_2 | 37218908 | 35111296 | 4.39G | 0.04 | 93.28 | 87.37 | 51.45 |
| Brown_1 | 27499268 | 26045606 | 3.26G | 0.04 | 94.99 | 89.99 | 51.51 |
| Brown_2 | 27499268 | 26045606 | 3.26G | 0.04 | 93.48 | 87.74 | 51.50 |

Raw reads: the numbers of original data; Clean reads: the numbers of after filtering the original
data; Error rate: Phred score, Qphred = $-10\log_{10}(e)$ ; Q20、Q30: the percent of the bases of
Qphred>20 or 30 in the toal bases.

**Figure 1.**
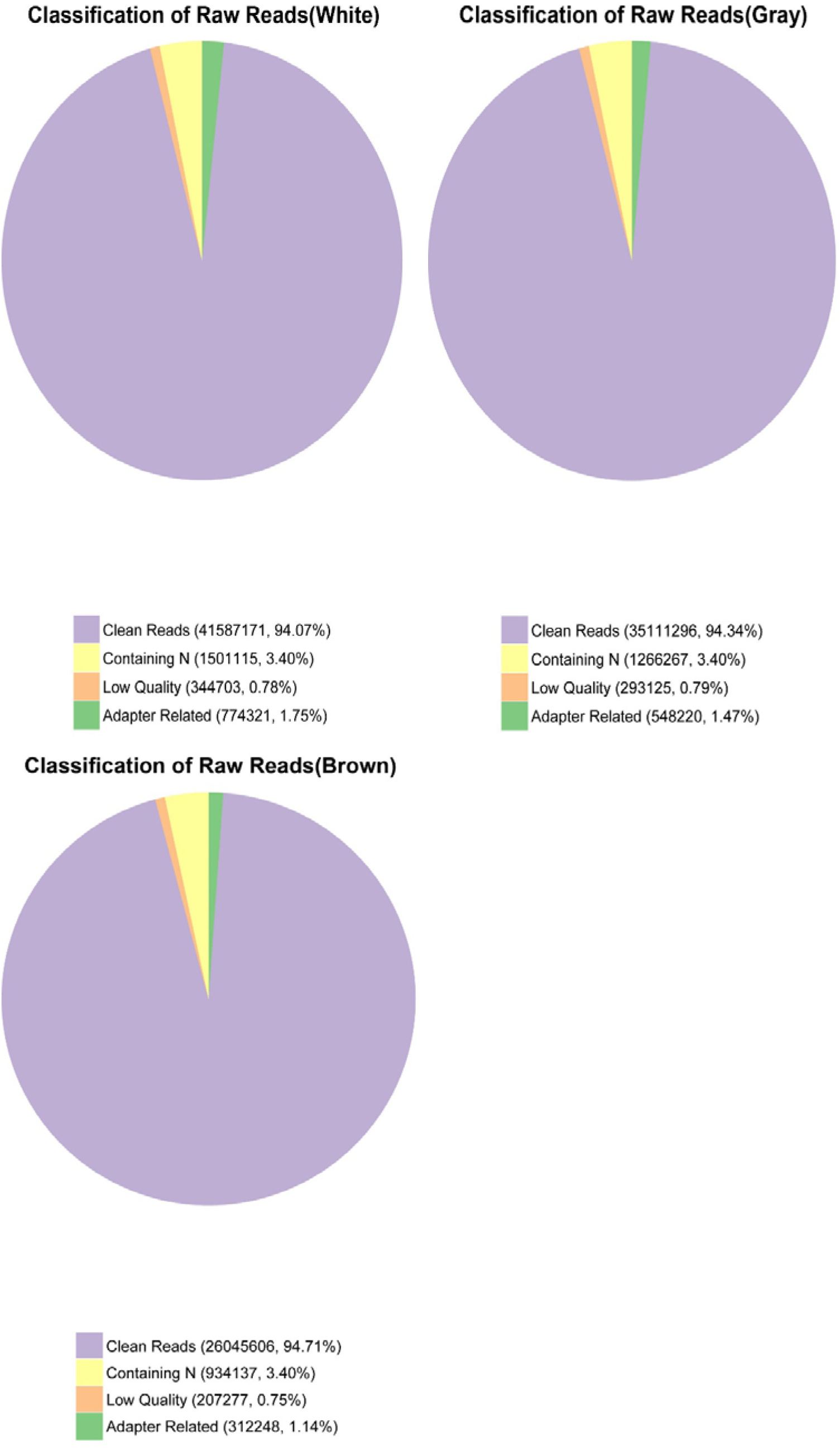
Classification of total raw reads pairs of different coat colors goat skin. After filtering the only adaptor sequences, containing N sequences and low quality sequences, the three RNA-Seq libraries still generated over 26 million clean reads pairs in each library, and the percentage of clean reads among raw tags in each library ranged from 94.07% to 94.71%.

Of the total reads, more than 77% matched either to a unique or to multiple genomic locations; the remaining were unmatched (Table 3), because only reads aligning entirely inside exonic regions will be matched (reads from exon-exon junction regions will not be matched). Of the total mapped, more than 78% matched to exon and more than 11% matched to intron; the remaning were matched the intergenic (Figure 2).

**Table 3.** Summary of read numbers based on the RNA-Seq data from different coat colors goat skin.

| Sample_name | White | Gray | Brown |
| --- | --- | --- | --- |
| Total reads | 83174342 | 70222592 | 52091212 |
| Total mapped | 65032149 (78.19%) | 54279674 (77.3%) | 40999750 (78.71%) |
| Multiple mapped | 1350169 (1.62%) | 1123310 (1.6%) | 833097 (1.6%) |
| Uniquely mapped | 63681980 (76.56%) | 53156364 (75.7%) | 40166653 (77.11%) |
| Non-splice reads | 39477742 (47.46%) | 33330204 (47.46%) | 25547588 (49.04%) |
| Splice reads | 24204238 (29.1%) | 19826160 (28.23%) | 14619065 (28.06%) |

**Figure 2.**
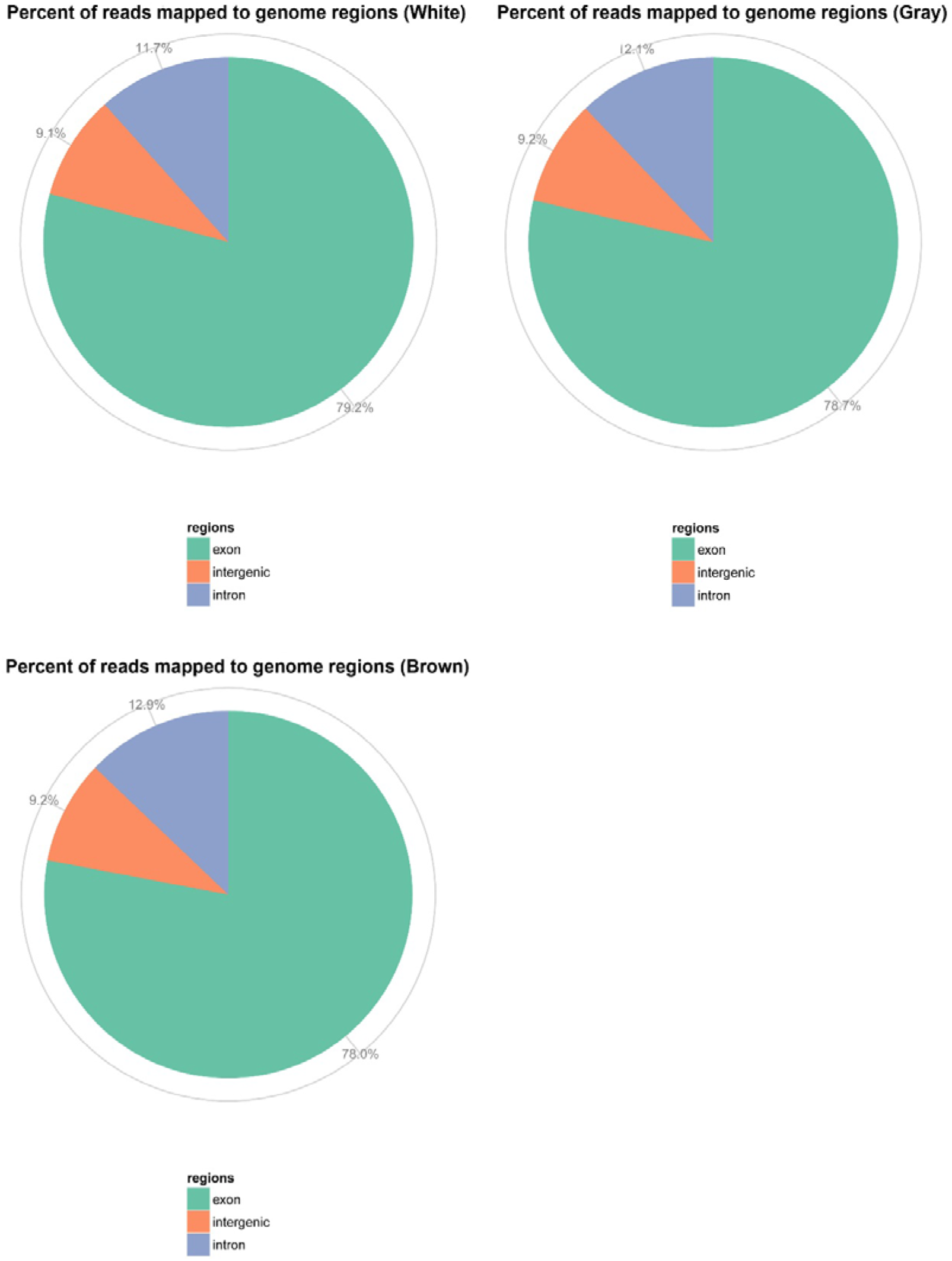
Percent of reads mapped to genome regions. The reads mapped to genome regions including exon, intron and intergenic.

### Genes highly expressed in goat skin

The top 30 genes most highly expressed in goat skin included genes of the keratin family and ribosomal proteins (Table S1). The most highly expressed gene in all three different coat colors goat skin was keratin associated protein 3-1. The other highly expressed genes in the different coat color goat skins included, trichohyalin, transcript variant X1, keratin 71, keratin 5, keratin 25, eukaryotic translation elongation factor 1 alpha 1, keratin 14, transcript variant X1, keratin 27, keratin, type II microfibrillar, component 7C-like, collagen, type I, alpha 1, uncharacterized LOC102168701, ribosomal protein L35, secreted protein, acidic, cysteine-rich (osteonectin), ribosomal protein L13a.The list of most highly expressed genes also included 2 unknown genes.

### Differentially expressed genes in different coat colors goat skin

To identify the differentially expressed genes (DEGs) of different coat colors goat skin, the differences in gene expression patterns were analysed for the pairs of the gray skin and white skin, the brown skin and white skin, the brown skin and gray skin. There were a total of 89 known genes and 2 novel genes identified as differentially expressed in gray skin versus white skin, of which 17 were down-regulated and 74 (including 2 novel genes) were up-regulated in skin from gray goat compared with skin from white goat (Figure 3 and Table S2). Between the brown and white libraries, 67 genes (including 1 novel gene) were differentially expressed, including 44 down-regulated (including 1 novel gene) and 23 up-regulated genes (Figure 3 and Table S3). When we compared the brown and gray libraries,154 DEGs (including 6 novel genes) were found, with 121 down-regulated (including 5 novel genes) and 33 up-regulated (including 1 novel gene) (Figure 3 and Table S4). This suggests that the differentiation of expressed genes between the brown skin and gray skin is larger than that between the gray skin and white skin, while a relatively smaller differentiation arises between the brown skin and white skin.

**Figure 3.**
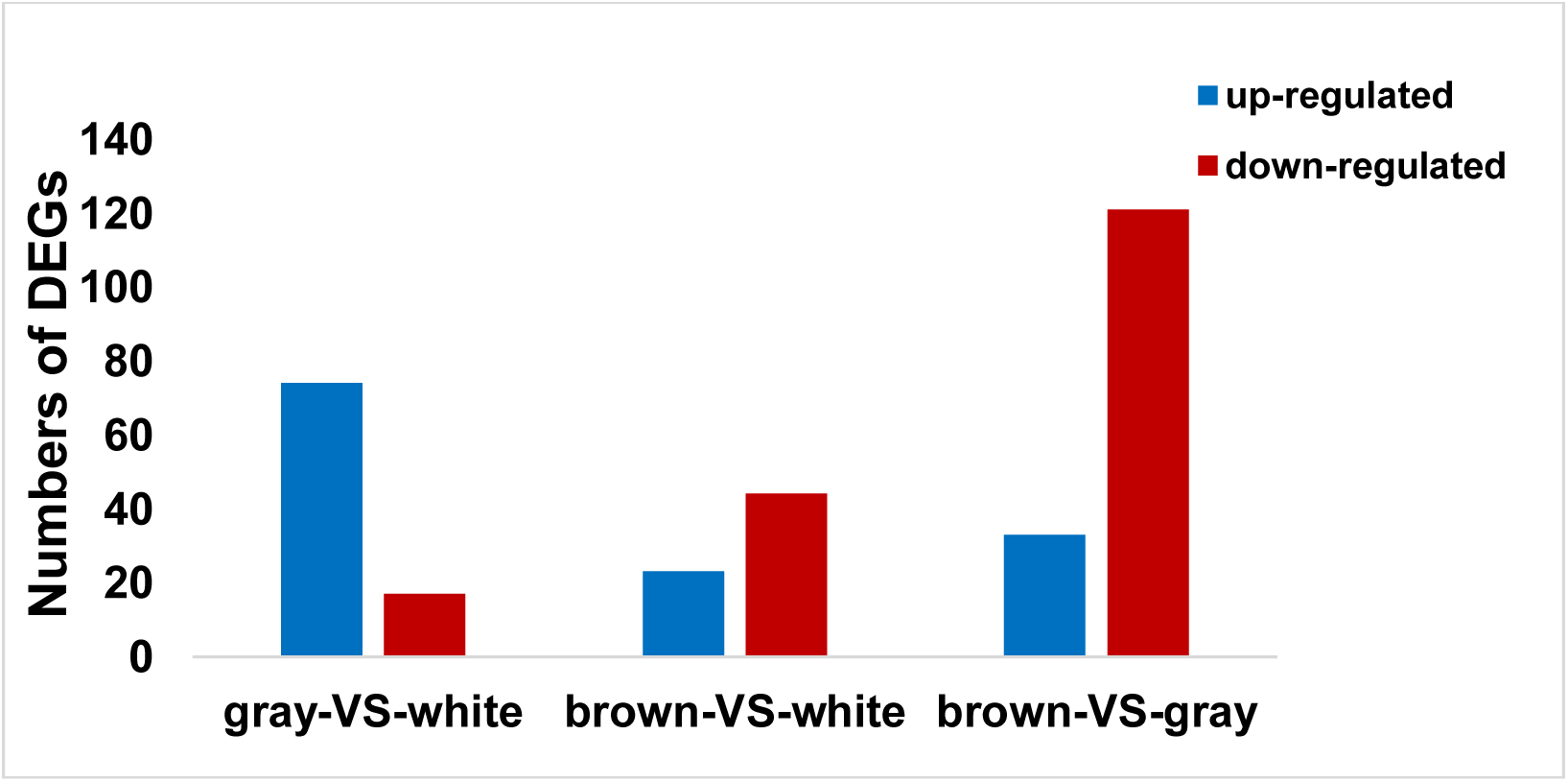
The numbers of DEGs between the three different coat colors goat skin. Between the gray skin and white skin libraries, there were 74 upregulated genes and 17 downregulated genes; Between the brown skin and white skin libraries, there were 23 upregulated genes and 44 downregulated genes, while there were 33 upregulated genes and 121 downregulated genes between the brown skin and gray skin libraries.

### Functional Classification Analysis

For the GO analysis of the gray versus white skin, 1249, 349 and 607 differentially expressed genes were grouped in biological process, cellular component and molecular function, respectively. Most of the differentially expressed genes were classified into two GO categories (single-organism metabolic process, oxidoreductase activity and catalytic activity; Figure 4 and Table S5). The majority of the GO terms including pigmentation do not appear to be significantly enriched in the differentially expressed genes. For the GO analysis of brown versus white skin, 505, 326 and 391 differentially expressed genes were grouped in biological process, cellular component and molecular function, respectively. Most of the differentially expressed genes were classified into three GO categories (oxidation-reduction process, extracellular region and structural molecule activity; Figure 4 and Table S5). The majority of the GO terms including pigmentation do not appear to be significantly enriched in the differentially expressed genes. For the GO analysis of brown versus white skin, 2368, 761 and 1100 differentially expressed genes were grouped in Most of the differentially expressed genes were classified into two GO categories (oxidation-reduction process, single-organism metabolic process and oxidoreductase activity; Figure 4 and Table S5). The majority of the GO differentially expressed genes.

**Figure 4.**
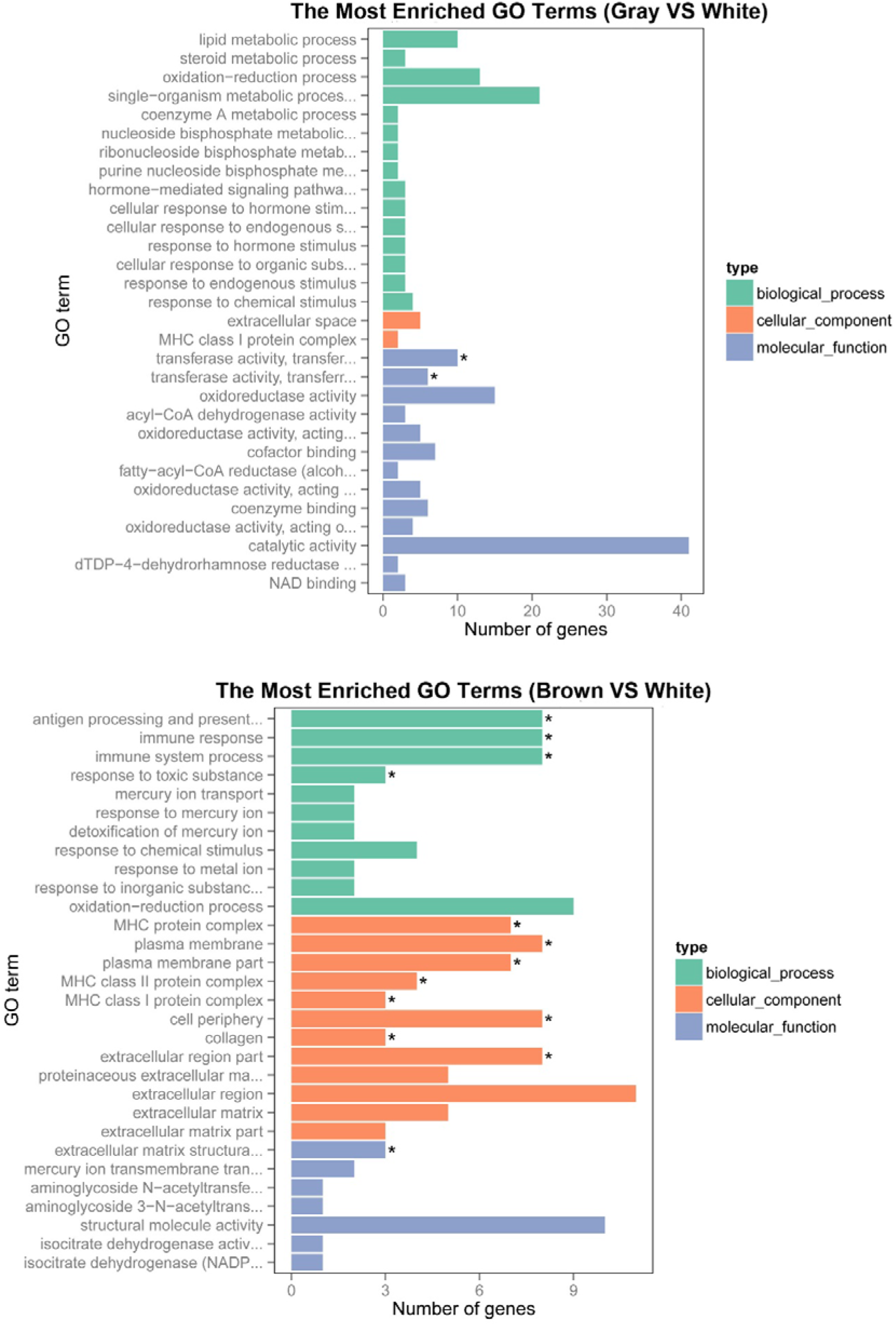
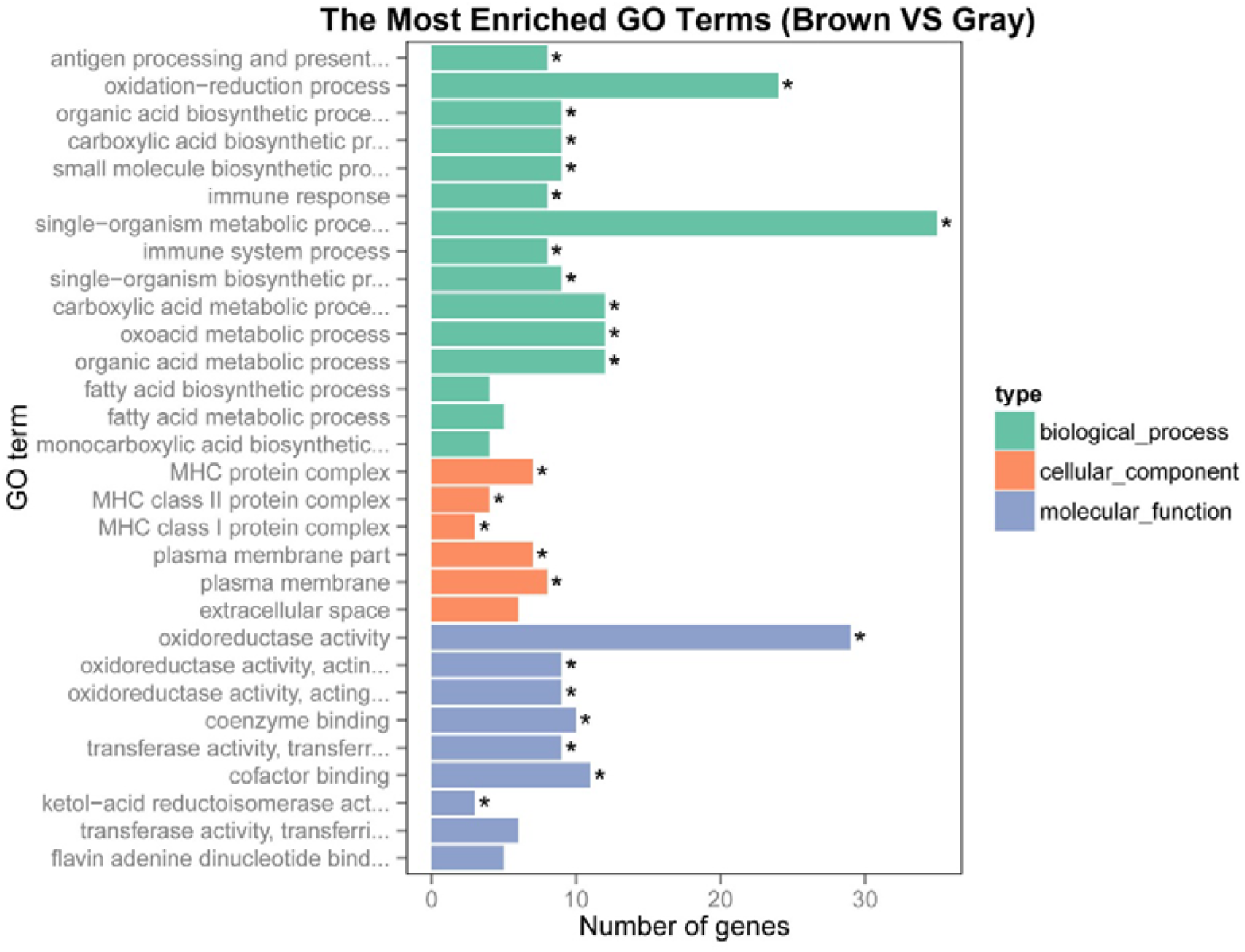
GO functional analysis of DEGs based on RNA-Seq data. The results were summarised the most enriched GO terms in three main categories: biological process, cellular component and molecular function.

In order to validate the transcriptome sequencing results, we selected 5 genes at random for real time PCR to determine their relative expression in different coat clor goat skin. These genes, identified as differentially expressed in dfferent coat color goat skin based on transcriptome sequencing analysis. The results of the qPCR were consistent with the RNA-seq except the expression of *TYRP1* between the gray skin and white skin libraries (Figure 5).

**Figure 5.**
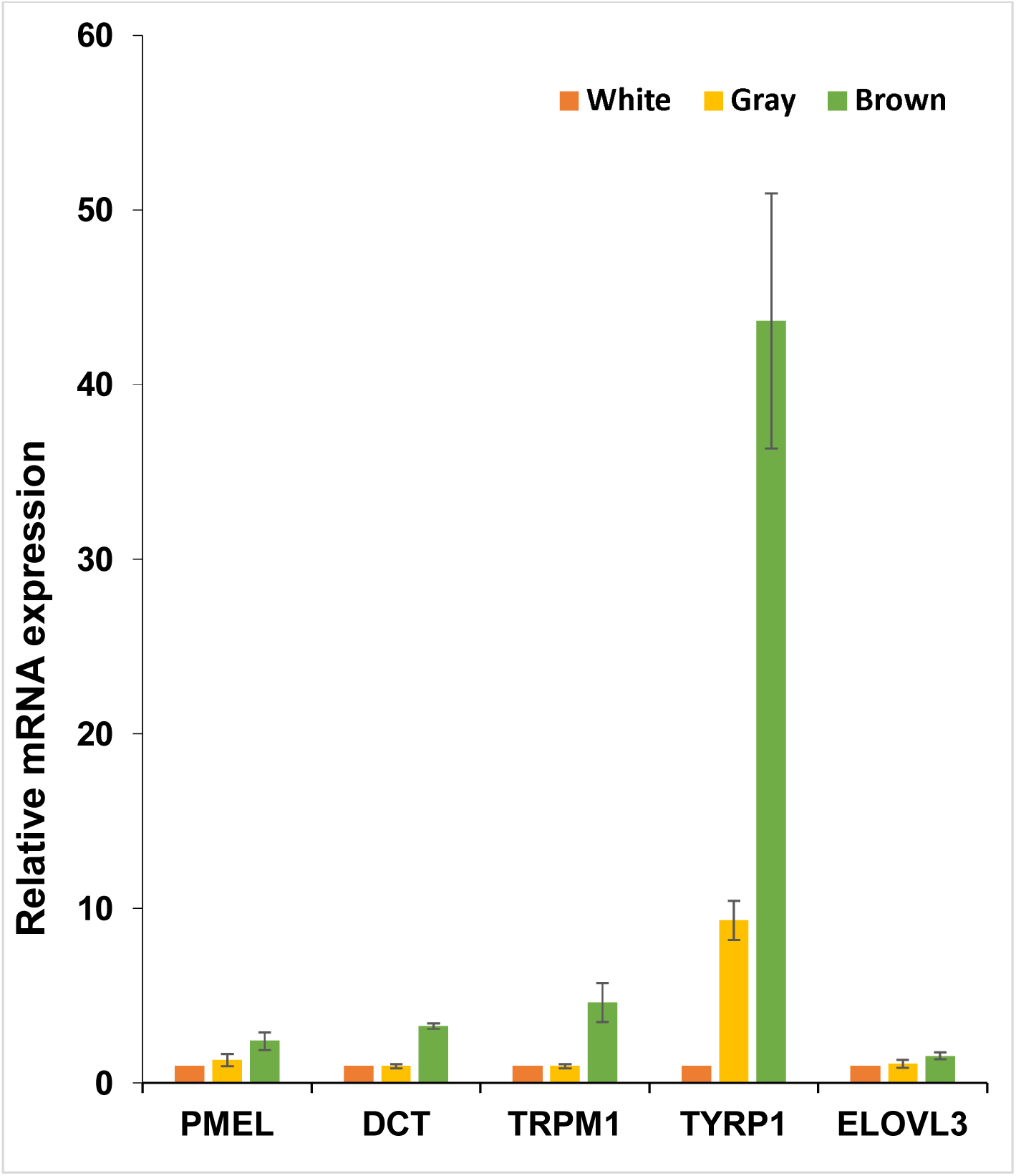
The qPCR validation of differentially expressed genes in the three different coat colors goat skin. Abundance of target genes was normalized relative to abundance of *GAPDH* gene. Bars in each panel represent the mean ± standard error (n = 3).

### KEGG pathway analysis

KEGG is a largely publicly available pathway-related database and is useful in searching for genes involved in metabolic or signal transduction pathways that were significantly enriched (Kanehisa *et al*. 2008). Of the 89 known genes differentially expressed in gray versus white goat skin, 40 had a specific KEGG pathway annotation. Of these KEGG pathway annotated genes, 3 were down-regulated in gray goat skin. These down-regulated genes are mainly involved in cysteine and methionine metabolism, melanogenesis, ribosome, metabolic pathways. Remaining KEGG pathway annotated genes were associated with 61 pathways including those functionally related to coat color in skin such as tyrosine metabolism (Table S6).

Between the brown and white goat skin, 23 DEGs with a KEGG pathway annotation were found and 14 were down-regulated in brown goat skin. These down-regulated genes were associated with 41 pathways including those functionally related to coat color in skin such as melanogenesis. Remaining KEGG pathway annotated genes were associated with 31 pathways including those functionally related to coat color in skin such as melanogenesis, tyrosine metabolism (Table S7).

Between the brown and gray goat skin, 72 DEGs with a KEGG pathway annotation were found and 63 were down-regulated in brown goat skin. These down-regulated genes were associated with 98 pathways including those functionally related to coat color in skin such as tyrosine metabolism. Remaining KEGG pathway annotated genes were associated with 29 pathways including those functionally related to coat color in skin such as melanogenesis, tyrosine metabolism (Table S8).

### Differential expression of known coat color genes

Approximately 171 cloned genes involved in different pathways controlling coat color formation have been identified in the mouse (http://www.espcr.org/micemut/#cloned). Those known coat color genes are routinely classified into six general functions including: Melanocyte development, Components of melanosomes and their precursors, Melanosome construction/protein routing, Melanosome transport, Eumelanin and pheomelanin and Systemic effects (Bennett and Lamoreux 2003). Expression of a total of 128 of aforementioned coat color genes was detected in goat skin in present studies, and 2 genes showed higher expression in gray goat skin versus white goat skin and 1 gene showed higher expression in white goat skin, 4 genes showed higher expression in brown goat skin versus white goat skin and 1 gene showed higher expression in white goat skin, 4 genes showed higher expression in brown goat skin versus gray goat skin. Interestingly, the expression of *ASIP* (agouti signaling protein) were almost not detected in the gray skin and the brown skin, but there was high expression in the white goat skin for *ASIP*. *PMEL*, *DCT*, *TRPM1* and *TYRP1* were significantly up-regulated in brown goat skin (brown vs white, brown vs gray), but *DCT*, *TRPM1* and *TYRP1* showed no differences in expression between the gray and white skin samples and only *PMEL* was significantly up-regulated in gray goat skin versus white goat skin. *ELOVL3* was significantly up-regulated in gray goat skin (gray vs white, brown vs gray). However, there were no significant differences in the expression of *ELOVL3* between the brown and white skin samples. *MC1R*, *MITF, TYR*, *KIT* and *KITLG* showed no differences when comparing the three different coat color groups. The genes associated with oculocutaneous albinism (OCA) such as *HPS1*, *HPS3*, *HPS4*, *HPS5* and *HPS6* were expressed in goat skin but all of them did not show differential expression associated with coat color.

## DISCUSSION

Mammalian coat color exhibits a wide range of shades and is dictated by melanin production in melanocytes (melanogenesis). Melanogenesis involves a complex molecular regulation (Slominski *et al*. 2004). In order to understand the molecular mechanisms of coat color formation, previous studies have reported the global gene expression profiles in skin of sheep and chicken with white versus black skin through Illumina sequencing technology (Fan *et al*. 2013; Zhang *et al*. 2015). Another study examined differences in gene expression associated with black spots infleece of Corriedale sheep using microarray technology (Penagaricano *et al*. 2012).

Our study offers new information related to gene expression profiles for three different coat colors goat skin. Our data analysis was based on the NCBI database of the goat reference genome (Capra hircus (assembly CHIR_1.0)).

To further investigate genes that may play important roles in goat skin, particularly in fiber/coat pigmentation, over 200 million transcriptome sequence reads were generated from different coat colors goat skin using high throughput RNA deep sequencing. From these reads there were 24,086 known genes identified as expressed in goat skin, among which 89 were differentially expressed in gray versus white goat skin, 66 were differentially expressed in brown versus white goat skin and 148 were differentially expressed in brown versus gray goat skin. It is acknowledged that study design was not optimal due to limited biological replication because single pooled samples (n=3 per coat color) were used in transcriptome sequencing analysis and the same three samples from different coat colors goat skin were used individually for quantitative real time PCR validation of the sequencing results. Despite such limitations, results have significantly enhanced understanding of goat skin transcriptome composition and potential differences in gene expression associated with coat color that are foundational to further study in the future.

Genes encoding for keratin family members and keratin associated proteins, ribosomal proteins were among the most highly expressed genes detected in goat skin. Of the top 30 highly expressed genes in goat skin, all 9 keratin family members and keratin associated proteins displayed no diffrenence between the three different coat colors goat skin. Hair keratins contain a much higher content of cysteine residues in their non-helical domains and thus form tougher and more durable structures via inter molecular disulfide bond formation (Mogosanu *et al*. 2014). Therefore, high expression of keratins is likely crucial for fleece strength. The ribosome is a central player in the translation system and its function is to decode the nucleotide sequence carried by the mRNA and convert it into anamino acid primary structure (Marshall *et al*. 2008). Abundant presence of ribosomal proteins in goat skin suggests the importance of high rates of protein translation in goat skin. In channel catfish skin, the expression of ribosomal proteins was high presumably due to higher levels of translational activities (Karsi *et al*. 2002; Patterson *et al*. 2003). It was also found that several collagen family members, particularly collagen, type I family members, were high expression in the three different coat colors goat skin. Collagen, in the form of elongated fibrils, is mostly found in fibrous tissues such as tendons, ligaments and skin (Zuber *et al*. 2015). Collagen has many medical uses in treating complications of the bones and skin. So high expression collagen may play important physiological function for skin. The expression of trichohyalin, transcript variant X1 was also high expression in the three different coat colors goat skin. Trichohyalin confers mechanical strength to the hair follicle inner root sheath and to other toughened epithelial tissues, such as the hard palate and filiform ridges of the tongue, by forming multiple complex crosslinks with itself and with other structural proteins (Steinert *et al*. 2003), therefore high expression trichohyalin plays important role in hair follicle development.

The GO and KEGG pathway analyses of differentially expressed genes revealed that most were associated with the function of single-organism metabolic process, oxidoreductase activity and oxidation-reduction process ontology categories. Of particular interest in our dataset were pathways related to pigmentation and melanogenesis. In this study, in terms of the molecular function category, 3 significant genes were found to be involved in melanogenesis and 3 significant genes were found to be involved in tyrosine metabolism. All these results provide strong evidence that there is a significant difference in the level of melanogenesis between the different coat colors goat skin. However, further investigation is still needed to confirm the regulatory relationships of these genes.

Melanocortin-1 receptor (*MC1R*) is responsible for binding melanocyte stimulating hormone (MSH), which is expressed by stressed keratinocytes, and initiating the cascade of melanogenesis (Solano *et al*. 2006). *MC1R* has classically been considered to play a role in and be the irreplaceable target involved in regulating mammalian skin pigmentation and hair color (Roberts *et al*. 2006; Schaffler *et al*. 2006; Lalueza-Fox *et al*. 2007). Recently, a study found that there is no association between *MC1R* polymorphisms and plumagecoloration in the zebra finch (Hoffman *et al*. 2014) which is consistent with another study that was performed in leaf warblers (Macdougall-Shackleton *et al*. 2003). Another recently study also indicated that the coat color variation observed in rhesusmacaques (variation from light to dark) is unlikely to be due to differences in the expression levels of the *MC1R* gene. In a recent microarray experiment, no significant differences in the expression of *MC1R* were observed between the white skin and black spots of sheep (Penagaricano *et al*. 2012). There were no significant differences in the expression of *MC1R* between the black-skinned and white-skinned Lueyang chickens (Zhang *et al*. 2015). Similarly and to our surprise, *MC1R* showed no significant differences in expression between the three different coat colors goat used in this study. The specific function and role of *MC1R* in pigmentation still requires further research.

The gene for agouti signaling protein (*ASIP*) is centrally involved in the expression of coat-color traits in animals. Previous studies have shown that the *ASIP* gene is responsible for the skin color of both white and black-coated sheep (Norris and Whan 2008) and the *ASIP* CNV were association with coat color in Tibetan sheep (Han *et al*. 2015). A mutation in *ASIP* causes the black-and-tan pigmentation phenotype observed pigs (Drogemuller *et al*. 2006), A missense mutation in the agouti signaling protein gene *(ASIP*) is associated with the no light points coat phenotype in donkeys (Abitbol *et al*. 2015). In our study, *ASIP* was found to show significantly different expression levels in the gray-skinned goats versus the white-skinned goats and the brown-skinned goats versus the white skinned goats, while *ASIP* was only detected in the white skin and almostly not detected in the gray skin and brown skin. The result that *ASIP* only was detected in the white skin and not in the dark skins seems to be inconsistent with the previous study, which has shown that the expression of the *ASIP* gene was also significantly greater in the black-skinned compared to the white-skinned chickens (Zhang *et al*. 2015). *ASIP* may play important role in the pigmentation of goat and further investigations are still needed to confirm the mechanism of *ASIP*.

*TYRP1*, one of themembers of the tyrosinase family, is a I type membrane bound protein that is expressed in both melanocytes and the retinal epithelium. *TYRP1* is involved in the distal eumelanic pathway and plays a role in stabilizing TYR, which is the critical and rate-determining enzyme in melanogenesis (Kobayashi *et al*. 1998). There existed a significant association between coat color and *TYRP1* in Soay sheep. In the free-living Soay sheep, coat color is either dark brown or light tawny color. The light phenotype is determined by homozygosity of a single recessive amino acid change (G→T transversion) at coding position 869 in the *TYRP1* gene (Gratten *et al*. 2007). This is consistent with studies in domestic sheep, where light coat color is known to be due to a decrease in the ratio of eumelanin to pheomelanin, relative to black coat color (Aliev *et al*. 1990). In Bionda dell’Adamello goat, the genotype frequency distribution of a non-synonymous SNP (G1112A) suggested a possible role of *TYRP1* in brown eumelanic goat coat colour (Nicoloso *et al*. 2012). The brown coat colour of Coppernecked goats is associated with a non-synonymous variant (p.Gly496Asp) at the *TYRP1* locus (Becker *et al*. 2015). In our study, *TYRP1* were significantly up-regulated in brown goat skin (brown vs white, brown vs gray) and *TYRP1* showed no differences in expression between the gray and white skin samples using high throughput RNA deep sequencing, but the subsequent validation were inconsistent with our transcriptome sequencing data. The results of the qPCR showed that *TYRP1* were significantly up-regulated in brown goat skin (brown vs white, brown vs gray) and had significantly higher expression in gary goat skin compared with white goat skin.These results still require additional research.

*PMEL* (*SILV* or *PMEL17*) is involved in melanosomal structure and acts as a scaffold in melanosomes by forming the proteolytic fibrillar matrix where melanin is deposited (Theos *et al*. 2005). It is a prominent member of the group of dilution genes. *SILV* mutations, which result in pigment dilution, have been discovered in many animals (Kerje *et al*. 2004; Brunberg *et al*. 2006; Clark *et al*. 2006; Gutierrez-Gil *et al*. 2007; Kuhn and Weikard 2007; Hellstrom *et al*. 2011). In our study, *PMEL* were significantly up-regulated in brown goat skin (brown vs white, brown vs gray) and had significantly higher expression in gary goat skin compared with white goat skin. The results of our RNA-seq and the subsequent validation were consistent.

*DCT* (*TYRP2*) is a melanogenic enzyme termed DOPAchrome tautomerase, which isomerizes the pigmented intermediate DOPAchrome to DHICA (5,6-dihydroxyindole-2-carboxylic acid) rather than to DHI (5,6-dihydroxyindole) (Tsukamoto *et al*. 1992). Mutations of *DCT* were associated with coat-color variation in mice (Jackson *et al*. 1992; Budd and Jackson 1995). The mice deficient in *DCT* showed a diluted coat color phenotype (Guyonneau *et al*. 2004). Mutations in dopachrome tautomerase (*Dct*) affect eumelanin/pheomelanin synthesis (Costin *et al*. 2005). Sheep with the GG genotype of *DCT* had significantly (P < 0.05) lower tyrosinase activity, alkali-soluble melanin content, and ratio of eumelanin: total melanin than sheep with GA and AA genotypes when measured across all investigated samples (Deng *et al*. 2009). *DCT* did not show expression in duck feather bulbs but expressed in retina (Li *et al*. 2012). Our RNA-seq and qPCR all showed that *DCT* were significantly up-regulated in brown goat skin (brown vs white, brown vs gray) and there were no significant differences in the expression of *DCT* between the gray and white skin samples.

*TRPM1* is a member of the transient receptor potential (TRP) channel family. Although the potential effects of *TRPM1* on pigmentation have not been fully elucidated, it has been reported to be a potent regulator of pigmentation (Oancea *et al*. 2009). Moreover, it is associated with a distinct leucistic white phenotype, the so-called leopard complex spotting, which is found in some horse breeds (Bellone *et al*. 2008; Wade *et al*. 2009). Interestingly, this mutation is linked with night blindness in homozygous Appaloosa horses. In homozygous leopard spotting horses, mRNA expression of *TRPM1* is significantly downregulated in both skin and retina (Bellone *et al*. 2008; Bellone *et al*. 2010). This gene is also involved in night blindness in humans (Van Genderen *et al*. 2009; Koike *et al*. 2010). Our RNA-seq and qPCR all showed that *TRPM1* were significantly up-regulated in brown goat skin (brown vs white, brown vs gray) and there were no significant differences in the expression of *TRPM1* between the gray and white skin samples.

*ELOVL3* encodes a protein that belongs to the GNS1/SUR4 family. Members of this family play a role in elongation of long chain fatty acids to provide precursors for synthesis of sphingolipids and ceramides. The Elovl3-ablated mice displayed a sparse hair coat (Westerberg *et al*. 2004). Our RNA-seq and qPCR all showed that *ELOVL3* were significantly up-regulated in gray goat skin (gray vs white, brown vs gray) and there were no significant differences in the expression of *ELOVL3* between the brown and white skin samples.

*ASIP* was only detected in the white skin and not in the dark skins, therefore *ASIP* may play important role in maintaining goat white coat color. *PMEL,TYRP1, TRPM1* and *DCT* were significantly up-regulated in brown goat skin compare with the gray and white color skin, so they may play important role in the formation of goat brown coat color. In addition, *ELOVL3* were significantly up-regulated in gray goat skin compare with the brown and white color skin and there were no significant differences in the expression of *ELOVL3* between the brown and white skin samples. *PMEL* had significantly higher expression in gary goat skin compared with white goat skin and it was also found that *TYRP1* had significantly higher expression in gary goat skin compared with white goat skin by qPCR. *ELOVL3, PMEL* and *TYRP1* may be important for the goat gray coat color although the result of *TYRP1* qPCR were inconsistent with RNA-seq. *PMEL* may be a key factor in the pigmentation of the three different coat colors. These results still require additional research.

## CONCLUSIONS

In summary, this is an original report of the transcriptome analysis of the skin from three different coat colors goat. The present study described and revealed a set differentially expressed known and novel genes found in goat skin that are potentially related to skin color and other physiological functions. The *MC1R* gene showed no difference in expression between the three diferent coat colors and the *ASIP* gene was only detected in the white skin, which are of particular interest for future studies to elucidate its functional roles in the regulation of skin color. *PMEL,TYRP1, TRPM1* and *DCT* may play important role in the formation of goat brown coat color and *ELOVL3, PMEL* and *TYRP1* may be important for the goat gray coat color. Results are foundational for future studies to potentially manipulate coat color via pharmacological and genetic approaches.

## ACKNOWLEDGMENTS

We are grateful to members of our laboratories for critical reading of the manuscript and helpful discussion. This work was supported by the National Natural Science Foundation of China (no: 31172196, 31501940), the Applied Basic Research Programs of Hebei Province (no:15962901D), College Innovation Team Leader Training Programme of Hebei Province (no: LJRC004), the Science and Technology Pillar Program of Qinhuangdao (no: 201502A058) and PhD Research Start Fund of Hebei Normal University of Science and Technology (2014). RNA sequences were completed by Novogene Bioinformatics Institute.

## LITERATURE CITED

Abitbol, M., R. Legrand and L. Tiret, 2015 A missense mutation in the agouti signaling protein gene (ASIP) is associated with the no light points coat phenotype in donkeys. Genet Sel Evol 47: 28.

Aliev, G., M. Rachkovsky, S. Ito, K. Wakamatsu and A. Ivanov, 1990 Pigment types in selected color genotypes of Asiatic sheep. Pigment Cell Res 3: 177–180.

Becker, D., M. Otto, P. Ammann, I. Keller, C. Drogemuller et al., 2015 The brown coat colour of Coppernecked goats is associated with a non-synonymous variant at the TYRP1 locus on chromosome 8. Anim Genet 46: 50–54.

Bellone, R. R., S. A. Brooks, L. Sandmeyer, B. A. Murphy, G. Forsyth et al., 2008 Differential gene expression of TRPM1, the potential cause of congenital stationary night blindness and coat spotting patterns (LP) in the Appaloosa horse (Equus caballus). Genetics 179: 1861–1870.

Bellone, R. R., G. Forsyth, T. Leeb, S. Archer, S. Sigurdsson et al., 2010 Fine-mapping and mutation analysis of TRPM1: a candidate gene for leopard complex (LP) spotting and congenital stationary night blindness in horses. Brief Funct Genomics 9: 193–207.

Bennett, D. C., and M. L. Lamoreux, 2003 The color loci of mice--a genetic century. Pigment Cell Res 16: 333–344.

Brunberg, E., L. Andersson, G. Cothran, K. Sandberg, S. Mikko et al., 2006 A missense mutation in PMEL17 is associated with the Silver coat color in the horse. BMC Genet 7: 46.

Budd, P. S., and I. J. Jackson, 1995 Structure of the mouse tyrosinase-related protein-2/dopachrome tautomerase (Tyrp2/Dct) gene and sequence of two novel slaty alleles. Genomics 29: 35–43.

Bunge, R., D. L. Thomas, T. G. Nash and C. J. Lupton, 1996 Performance of hair breeds and prolific wool breeds of sheep in southern Illinois: wool production and fleece quality. J Anim Sci 74: 25–30.

Cieslak, M., M. Reissmann, M. Hofreiter and A. Ludwig, 2011 Colours of domestication. Biol Rev Camb Philos Soc 86: 885–899.

Clark, L. A., J. M. Wahl, C. A. Rees and K. E. Murphy, 2006 Retrotransposon insertion in SILV is responsible for merle patterning of the domestic dog. Proc Natl Acad Sci U S A 103: 1376–1381.

Costin, G. E., J. C. Valencia, K. Wakamatsu, S. Ito, F. Solano et al., 2005 Mutations in dopachrome tautomerase (Dct) affect eumelanin/pheomelanin synthesis, but do not affect intracellular trafficking of the mutant protein. Biochem J 391: 249–259.

Deng, W., Y. Tan, X. Wang, D. Xi, Y. He et al., 2009 Molecular cloning, sequence characteristics, and polymorphism analyses of the tyrosinase-related protein 2 / DOPAchrome tautomerase gene of black-boned sheep (Ovis aries). Genome 52: 1001–1011.

Dereure, O., 2001 Drug-induced skin pigmentation. Epidemiology, diagnosis and treatment. Am J Clin Dermatol 2: 253–262.

Drogemuller, C., A. Giese, F. Martins-Wess, S. Wiedemann, L. Andersson et al., 2006 The mutation causing the black-and-tan pigmentation phenotype of Mangalitza pigs maps to the porcine ASIP locus but does not affect its coding sequence. Mamm Genome 17: 58–66.

Ducrest, A. L., L. Keller and A. Roulin, 2008 Pleiotropy in the melanocortin system, coloration and behavioural syndromes. Trends Ecol Evol 23: 502–510.

Duffy, D. L., Z. Z. Zhao, R. A. Sturm, N. K. Hayward, N. G. Martin et al., 2010 Multiple pigmentation gene polymorphisms account for a substantial proportion of risk of cutaneous malignant melanoma. J Invest Dermatol 130: 520–528.

Emaresi, G., A. L. Ducrest, P. Bize, H. Richter, C. Simon et al., 2013 Pleiotropy in the melanocortin system: expression levels of this system are associated with melanogenesis and pigmentation in the tawny owl (Strix aluco). Mol Ecol 22: 4915–4930.

Fan, R., J. Xie, J. Bai, H. Wang, X. Tian et al., 2013 Skin transcriptome profiles associated with coat color in sheep. BMC Genomics 14: 389.

Fan, Y., P. Wang, W. Fu, T. Dong, C. Qi et al., 2014 Genome-wide association study for pigmentation traits in Chinese Holstein population. Anim Genet 45: 740–744.

Garcia-Gamez, E., A. Reverter, V. Whan, S. M. McWilliam, J. J. Arranz et al., 2011 Using regulatory and epistatic networks to extend the findings of a genome scan: identifying the gene drivers of pigmentation in merino sheep. PLoS One 6: e21158.

Gratten, J., D. Beraldi, B. V. Lowder, A. F. McRae, P. M. Visscher et al., 2007 Compelling evidence that a single nucleotide substitution in TYRP1 is responsible for coat-colour polymorphism in a free-living population of Soay sheep. Proc Biol Sci 274: 619–626.

Gutierrez-Gil, B., P. Wiener and J. L. Williams, 2007 Genetic effects on coat colour in cattle: dilution of eumelanin and phaeomelanin pigments in an F2-Backcross Charolais x Holstein population. BMC Genet 8: 56.

Guyonneau, L., F. Murisier, A. Rossier, A. Moulin and F. Beermann, 2004 Melanocytes and pigmentation are affected in dopachrome tautomerase knockout mice. Mol Cell Biol 24: 3396–3403.

Han, J. L., M. Yang, Y. J. Yue, T. T. Guo, J. B. Liu et al., 2015 Analysis of agouti signaling protein (ASIP) gene polymorphisms and association with coat color in Tibetan sheep (Ovis aries). Genet Mol Res 14: 1200–1209.

Hart, K. L., S. L. Kimura, V. Mushailov, Z. M. Budimlija, M. Prinz et al., 2013 Improved eye-and skin-color prediction based on 8 SNPs. Croat Med J 54: 248–256.

Hellstrom, A. R., B. Watt, S. S. Fard, D. Tenza, P. Mannstrom et al., 2011 Inactivation of Pmel alters melanosome shape but has only a subtle effect on visible pigmentation. PLoS Genet 7: e1002285.

Hoffman, J. I., E. T. Krause, K. Lehmann and O. Kruger, 2014 MC1R genotype and plumage colouration in the zebra finch (Taeniopygia guttata): population structure generates artefactual associations. PLoS One 9: e86519.

Ito, S., and K. Wakamatsu, 2008 Chemistry of mixed melanogenesis--pivotal roles of dopaquinone. Photochem Photobiol 84: 582–592.

Ito, S., K. Wakamatsu and H. Ozeki, 2000 Chemical analysis of melanins and its application to the study of the regulation of melanogenesis. Pigment Cell Res 13 Suppl 8: 103–109.

Jablonski, N. G., and G. Chaplin, 2010 Colloquium paper: human skin pigmentation as an adaptation to UV radiation. Proc Natl Acad Sci U S A 107 Suppl 2: 8962–8968.

Jackson, I. J., D. M. Chambers, K. Tsukamoto, N. G. Copeland, D. J. Gilbert et al., 1992 A second tyrosinase-related protein, TRP-2, maps to and is mutated at the mouse slaty locus. EMBO J 11: 527–535.

Kanehisa, M., M. Araki, S. Goto, M. Hattori, M. Hirakawa et al., 2008 KEGG for linking genomes to life and the environment. Nucleic Acids Res 36: D480–484.

Karsi, A., D. Cao, P. Li, A. Patterson, A. Kocabas et al., 2002 Transcriptome analysis of channel catfish (Ictalurus punctatus): initial analysis of gene expression and microsatellite-containing cDNAs in the skin. Gene 285: 157–168.

Kerje, S., P. Sharma, U. Gunnarsson, H. Kim, S. Bagchi et al., 2004 The Dominant white, Dun and Smoky color variants in chicken are associated with insertion/deletion polymorphisms in the PMEL17 gene. Genetics 168: 1507–1518.

Kidson, S. H., and B. C. Fabian, 1981 The effect of temperature on tyrosinase activity in Himalayan mouse skin. J Exp Zool 215: 91–97.

Kobayashi, T., G. Imokawa, D. C. Bennett and V. J. Hearing, 1998 Tyrosinase stabilization by Tyrp1 (the brown locus protein). J Biol Chem 273: 31801–31805.

Koike, C., T. Numata, H. Ueda, Y. Mori and T. Furukawa, 2010 TRPM1: a vertebrate TRP channel responsible for retinal ON bipolar function. Cell Calcium 48: 95–101.

Kuhn, C., and R. Weikard, 2007 An investigation into the genetic background of coat colour dilution in a Charolais x German Holstein F2 resource population. Anim Genet 38: 109–113.

Lalueza-Fox, C., H. Rompler, D. Caramelli, C. Staubert, G. Catalano et al., 2007 A melanocortin 1 receptor allele suggests varying pigmentation among Neanderthals. Science 318: 1453–1455.

Lamoreux, M. L., K. Wakamatsu and S. Ito, 2001 Interaction of major coat color gene functions in mice as studied by chemical analysis of eumelanin and pheomelanin. Pigment Cell Res 14: 23–31.

Li, M. H., T. Tiirikka and J. Kantanen, 2014 A genome-wide scan study identifies a single nucleotide substitution in ASIP associated with white versus non-white coat-colour variation in sheep (Ovis aries). Heredity (Edinb) 112: 122–131.

Li, S., C. Wang, W. Yu, S. Zhao and Y. Gong, 2012 Identification of genes related to white and black plumage formation by RNA-Seq from white and black feather bulbs in ducks. PLoS One 7: e36592.

MacDougall-Shackleton, E. A., L. Blanchard, S. A. Igdoura and H. L. Gibbs, 2003 Unmelanized plumage patterns in Old World leaf warblers do not correspond to sequence variation at the melanocortin-1 receptor locus (MC1R). Mol Biol Evol 20: 1675–1681.

Marshall, R. A., C. E. Aitken, M. Dorywalska and J. D. Puglisi, 2008 Translation at the single-molecule level. Annu Rev Biochem 77: 177–203.

Minvielle, F., B. Bed'hom, J. L. Coville, S. Ito, M. Inoue-Murayama et al., 2010 The “silver” Japanese quail and the MITF gene: causal mutation, associated traits and homology with the "blue" chicken plumage. BMC Genet 11: 15.

Mogosanu, G. D., A. M. Grumezescu and M. C. Chifiriuc, 2014 Keratin-based biomaterials for biomedical applications. Curr Drug Targets 15: 518–530.

Nan, H., P. Kraft, D. J. Hunter and J. Han, 2009 Genetic variants in pigmentation genes, pigmentary phenotypes, and risk of skin cancer in Caucasians. Int J Cancer 125: 909–917.

Nicoloso, L., R. Negrini, P. Ajmone-Marsan and P. Crepaldi, 2012 On the way to functional agro biodiversity: coat colour gene variability in goats. Animal 6: 41–49.

Norris, B. J., and V. A. Whan, 2008 A gene duplication affecting expression of the ovine ASIP gene is responsible for white and black sheep. Genome Res 18: 1282–1293.

Oancea, E., J. Vriens, S. Brauchi, J. Jun, I. Splawski et al., 2009 TRPM1 forms ion channels associated with melanin content in melanocytes. Sci Signal 2: ra21.

Ozsolak, F., and P. M. Milos, 2011 RNA sequencing: advances, challenges and opportunities. Nat Rev Genet 12: 87–98.

Patterson, A., A. Karsi, J. Feng and Z. Liu, 2003 Translational machinery of channel catfish: II. Complementary DNA and expression of the complete set of 47 60S ribosomal proteins. Gene 305: 151–160.

Penagaricano, F., P. Zorrilla, H. Naya, C. Robello and J. I. Urioste, 2012 Gene expression analysis identifies new candidate genes associated with the development of black skin spots in Corriedale sheep. J Appl Genet 53: 99–106.

Roberts, D. W., R. A. Newton, K. A. Beaumont, J. Helen Leonard and R. A. Sturm, 2006 Quantitative analysis of MC1R gene expression in human skin cell cultures. Pigment Cell Res 19: 76–89.

Schaffler, A., J. Scholmerich and C. Buechler, 2006 The role of 'adipotropins' and the clinical importance of a potential hypothalamic-pituitary-adipose axis. Nat Clin Pract Endocrinol Metab 2: 374–383.

Slominski, A., D. J. Tobin, S. Shibahara and J. Wortsman, 2004 Melanin pigmentation in mammalian skin and its hormonal regulation. Physiol Rev 84: 1155–1228.

Solano, F., S. Briganti, M. Picardo and G. Ghanem, 2006 Hypopigmenting agents: an updated review on biological, chemical and clinical aspects. Pigment Cell Res 19: 550–571.

Steinert, P. M., D. A. Parry and L. N. Marekov, 2003 Trichohyalin mechanically strengthens the hair follicle: multiple cross-bridging roles in the inner root shealth. J Biol Chem 278: 41409–41419.

Steingrimsson, E., N. G. Copeland and N. A. Jenkins, 2006 Mouse coat color mutations: from fancy mice to functional genomics. Dev Dyn 235: 2401–2411.

Sturm, R. A., and D. L. Duffy, 2012 Human pigmentation genes under environmental selection. Genome Biol 13: 248.

Sulem, P., D. F. Gudbjartsson, S. N. Stacey, A. Helgason, T. Rafnar et al., 2007 Genetic determinants of hair, eye and skin pigmentation in Europeans. Nat Genet 39: 1443–1452.

Theos, A. C., S. T. Truschel, G. Raposo and M. S. Marks, 2005 The Silver locus product Pmel17/gp100/Silv/ME20: controversial in name and in function. Pigment Cell Res 18: 322–336.

Tsukamoto, K., I. J. Jackson, K. Urabe, P. M. Montague and V. J. Hearing, 1992 A second tyrosinase-related protein, TRP-2, is a melanogenic enzyme termed DOPAchrome tautomerase. EMBO J 11: 519–526.

Vage, D. I., H. Klungland, D. Lu and R. D. Cone, 1999 Molecular and pharmacological characterization of dominant black coat color in sheep. Mamm Genome 10: 39–43.

van Genderen, M. M., M. M. Bijveld, Y. B. Claassen, R. J. Florijn, J. N. Pearring et al., 2009 Mutations in TRPM1 are a common cause of complete congenital stationary night blindness. Am J Hum Genet 85: 730–736.

Wade, C. M., E. Giulotto, S. Sigurdsson, M. Zoli, S. Gnerre et al., 2009 Genome sequence, comparative analysis, and population genetics of the domestic horse. Science 326: 865–867.

Wang, L., J. Fan, M. Yu, S. Zheng and Y. Zhao, 2011a Association of goat (Capra hircus) CD4 gene exon 6 polymorphisms with ability of sperm internalizing exogenous DNA. Mol Biol Rep 38: 1621–1628.

Wang, P. Q., L. M. Deng, B. Y. Zhang, M. X. Chu and J. Z. Hou, 2011b Polymorphisms of the cocaine-amphetamine-regulated transcript (CART) gene and their association with reproductive traits in Chinese goats. Genet Mol Res 10: 731–738.

Wang, Z., M. Gerstein and M. Snyder, 2009 RNA-Seq: a revolutionary tool for transcriptomics. Nat Rev Genet 10: 57–63.

Westerberg, R., P. Tvrdik, A. B. Unden, J. E. Mansson, L. Norlen et al., 2004 Role for ELOVL3 and fatty acid chain length in development of hair and skin function. J Biol Chem 279: 5621–5629.

Wilhelm, B. T., and J. R. Landry, 2009 RNA-Seq-quantitative measurement of expression through massively parallel RNA-sequencing. Methods 48: 249–257.

Xu, T., X. Guo, H. Wang, F. Hao, X. Du et al., 2013 Differential gene expression analysis between anagen and telogen of Capra hircus skin based on the de novo assembled transcriptome sequence. Gene 520: 30–38.

Zhang, J., F. Liu, J. Cao and X. Liu, 2015 Skin transcriptome profiles associated with skin color in chickens. PLoS One 10: e0127301.

Zuber, M., F. Zia, K. M. Zia, S. Tabasum, M. Salman et al., 2015 Collagen based polyurethanes-A review of recent advances and perspective. Int J Biol Macromol 80: 366–374.

